# Self-generated brain-wide spiking cascades govern replay dynamics in the hippocampus

**DOI:** 10.1101/2022.09.05.506667

**Authors:** Yifan Yang, David A. Leopold, Jeff H. Duyn, Xiao Liu

**Affiliations:** Department of Biomedical Engineering, The Pennsylvania State University, University Park, PA, 16802, USA; Institute for Computational and Data Sciences, The Pennsylvania State University, University Park, PA, 16802, USA; Neurophysiology Imaging Facility, National Institute of Mental Health, National Institute of Neurological. Disorders and Stroke, and National Eye Institute, National Institutes of Health, Bethesda, MD, 20892, USA; Section on Cognitive Neurophysiology and Imaging, Laboratory of Neuropsychology, National Institute of Mental Health, National Institutes of Health, Bethesda, MD, 20892, USA; Advanced MRI Section, Laboratory of Functional and Molecular Imaging, National Institute of Neurological Disorders and Stroke, National Institutes of Health, Bethesda, MD 20892, USA

**Author notes:** Corresponding Author: Xiao Liu, PhD, 431 Chemical and Biomedical Engineering Building, The Pennsylvania State University, University Park, PA 16802-4400. **Author Contributions: Y.Y.&X.L.** contributed to the conception, design of the work, and data analysis; **X.L.** also devoted the efforts to the supervision, project administration and funding acquisition; **Y.Y., D.A.L., J.H.D. & X.L.** contributed to data visualization, and writing the paper.

## Abstract

During states of behavioral quiescence, neurons in the hippocampus replay sequences of spiking activity experienced in earlier behavioral episodes. While such replay sequences are hypothesized to serve learning and memory by facilitating synaptic consolidation, their generative mechanisms remain poorly understood. Increasing evidence suggests that they might be generated internally, or at least strongly constrained by internal circuit dynamics. Recent work demonstrated that, across the forebrain, approximately 70% of neurons participate in a pattern of sequential spiking cascades during rest. Like hippocampal replay sequences, these brain-wide spiking cascades occur together with high-frequency hippocampal ripples and therefore may share a common generative mechanism. Here we systematically investigated the relationship between replay activity and sequential spiking cascades by analyzing a database of intracortical electrocortical recordings in mice. For neuronal subpopulations in the hippocampus and visual cortex, we assessed spiking sequences elicited during video viewing as well as potential replay events during subsequent periods of rest. We found that replay events were unique to hippocampal time-sensitive neurons and occurred together with spiking cascades throughout the forebrain. Furthermore, forward and time-reversed replay sequences were associated with different types of spiking cascades. Overall, these findings indicate that hippocampal replay events are generated and structured according to resting state circuit dynamics manifest across a large portion of the brain.

## Introduction

Learning and memory are the cornerstone of intelligence. The hippocampus is a key brain structure involved in these functions. A remarkable finding about the rodent hippocampus is the fact that its place selective neurons (“place cells”) can replay sequences of activity previously induced by active exploration of a spatial environment. Often these replay episodes take place during rest and sleep and are typically manifest in a temporally compressed form (1–4). Such replay sequences are associated with prototypical electrical events originating in the hippocampus called sharp wave ripple complexes (SPW-R) (5–7). These events, which are evident as high-amplitude bursts in hippocampal local field recordings, have been proposed to play an important role in the consolidation of episodic memory (8–11).

The nature of hippocampal firing sequences during quiescent periods is a matter of active research. Increasing evidence suggests that these sequences, rather than being induced by the external experience itself, are fundamentally a product of internal circuit dynamics (12–16). A puzzling finding is that, in addition to replaying previous sequences generated during active behaviors, hippocampal place cells also appear to “preplay” a firing sequence that is only encountered later during exposure to a novel environment (17–19), suggesting that the hippocampal sequences exist before experience. Similarly, in a related subclass of hippocampal neurons, firing sequences are appear generated not by the registration of external events, but instead by the passage of time (20). The sequential firing of these “time neurons” can occur in the absence of changing environmental or body-derived inputs (21–24). These findings of preplays and time neurons have propelled a new theory that episode-specific activity sequences of hippocampal neuronal assembly roll forward as a result of self-organization of the brain and this temporal flow of activity is determined by intrinsic neuronal architecture (12–14).

A distinct type of sequential activations have recently been shown to shape neural firing across the forebrain beyond the hippocampus, across multiple cortical and thalamic structures (25). Like the replay, preplay, and time sequences observed in the hippocampus, these widespread patterns operate autonomously in the absence of external perturbations. They are expressed as stereotypic spiking cascades that affect a large proportion (∼70%) of the neural population in all tested forebrain areas. They are synchronized and quasi-periodic, with individual sequences lasting between 5-15 s. Moreover, each individual neuron bears a consistent temporal signature in its peak firing, leading or lagging the population peak by a fixed temporal interval. Importantly, these single-cell spiking sequences, which are expressed at many locations across the forebrain, were found to be synchronized with the slow modulation of hippocampal SPW-R occurrence (25). This synchronization with hippocampal ripples raises the question whether these widespread sequential spiking cascades might stem from the same self-generated brain dynamics as the hippocampal replays, which also concur with the hippocampal ripples as sequential activations.

In the present study, we investigated this possibility by analyzing population neuronal recordings from the visual cortex and hippocampus of the mouse under conditions conducive to replay activity. Using data available through the Allen Visual Coding project (26, 27), we first evaluated the activity of individual neurons in the visual cortex and hippocampus recorded during the viewing of a movie. Neurons in both areas yielded responses associated with particular moments or events in the movie, forming temporal sequences of neuronal spiking. The activity of these apparently time-selective neurons during subsequent periods of rest recapitulated the movie-induced sequence in a temporally compressed manner in the hippocampus, but not the visual cortex. We then investigated the relationship between these movie-related replay events and previously reported spontaneous firing cascades that engulf the brain during rest (25).

Importantly, the hippocampal replay events were temporally aligned to the spiking cascades, indicating that the replay activity in the hippocampus is one facet of a larger-scale pattern of sequential neural dynamics expressed spontaneously across the brain. A fine-scale analysis further revealed that forward and reverse hippocampal replays appeared respectively during two fundamental types of spiking cascade events of shorter duration. Together, these findings indicate that the hippocampal replay events are generated and structured according to resting state circuit dynamics manifest as the spiking cascade across a large portion of the brain.

## Results

We analyzed large-scale neuronal recordings in mice from the Visual Coding project of the Allen Institute. The dataset includes spiking activity of a large group of neurons simultaneously recorded from various brain cortical and subcortical regions. We focused on the spiking data of ∼10,000 neurons recorded from 14 mice in 44 brain regions (730 ± 178 neurons per mouse, mean ± SD) during two movie sessions and a spontaneous session (**Fig. 1A-1B**). In each of the two movie sessions (i.e., the pre-rest and post-rest ones), the same 30-sec movie clip was repeatedly presented to mice 30 times. The spontaneous session is free of visual stimulation, and the 14 mice remained stationary for extended periods of time (>20 minutes) (see Materials and Methods for stationary quantification).

**Figure 1.**
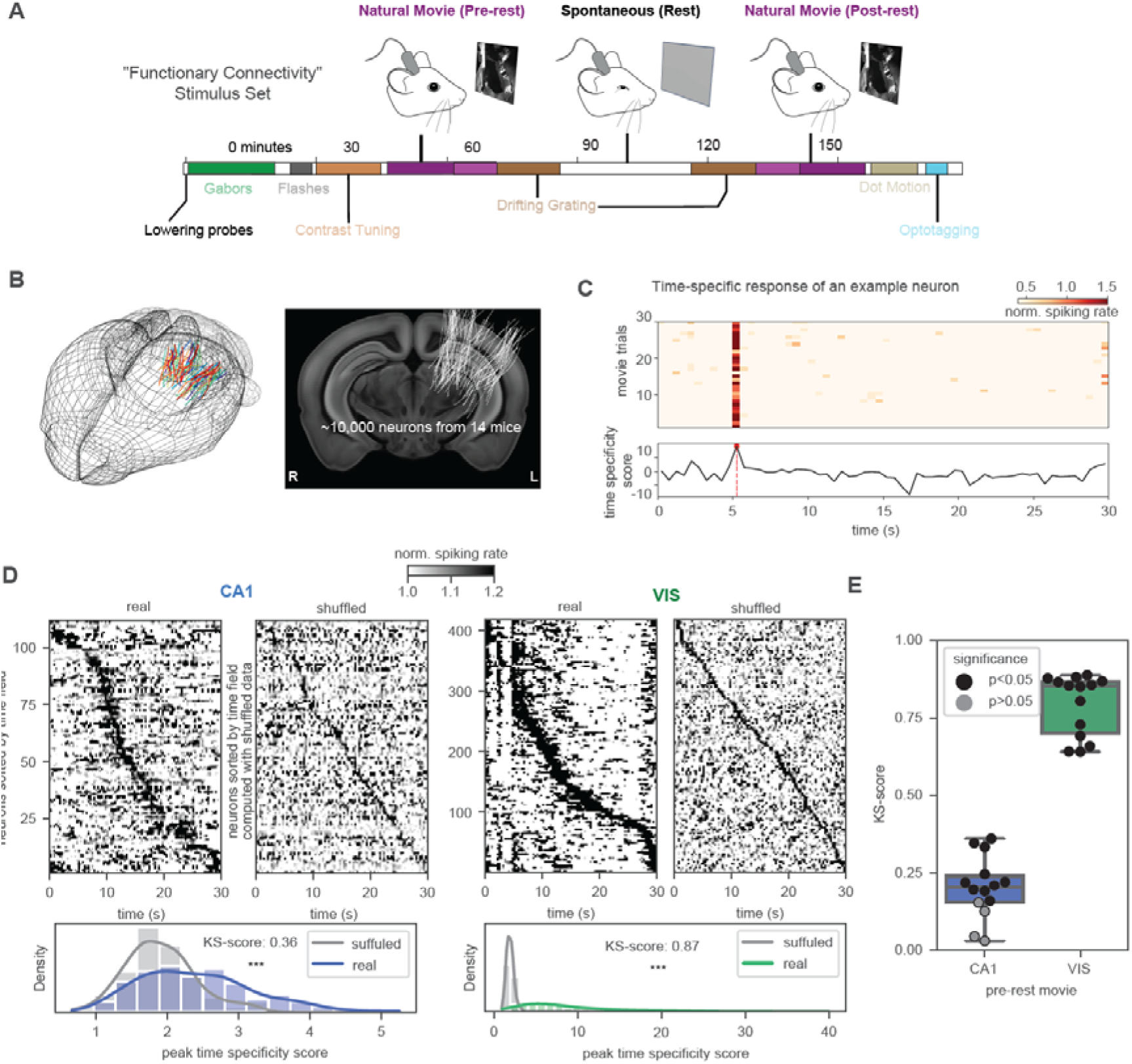
Time-specific responses of the hippocampal and visual neurons. (**A**) Illustration of the “Functional Connectivity” stimulus set of the “Visual Coding – Neuropixels” project, which includes a 30-minute spontaneous resting session and two natural movie sessions used in this study. (**B**) The three-dimensional (3D) location of 6,171 channels on 79 probes from 14 mice (left) and their projection onto the 2D middle slice of the brain template (right) in Allen Mouse Common Coordinate Framework. (**C**) An example neuron showing strong time-selective responses that are consistent across different trials of movie watching (top). The spiking rate was normalized to percentage changes with respect to its temporal mean. A time course of the time specificity score (bottom) achieved the peak value at the time field of this neuron (red line). (D) The averaged (N = 30 trials) spiking activity of the CA1 (left) and visual (right) neurons during the pre-rest movie watching. The neurons were sorted according to their time field. The two panels for each region show the results from the original data (left) and the shuffled control (r ght), which shuffled the spiking data of 0.5-sec time bins randomly within each movie trial. Distributions of the peak time specificity scores are compared between the real and shuffled data for both the CA1 (bottom left) and visual neurons (bottom right). The box plot of Kolmogorov–Smirnov (KS) score for all 14 mice. The KS score measures the difference in peak time specificity score distribution between the real and shuffled data. They are summarized for CA1 neurons and VIS neurons respectively. Each dot represents a mouse, and the black dot indicates a significant difference between the real and shuffled data (p<0.05).

### Hippocampal and visual neurons showed time-selective response during movie watching

We first examined time-selective responses of neurons in the hippocampus and the visual cortex during movie watching. To do this, a time course of time specificity score was computed for each neuron to quantify its firing rate increase at a specific moment (i.e., a 0.5-sec time bin) compared with other periods of the movie. The peak score quantifies the amplitude of the time-selective response, whereas the time to achieve this peak is regarded as the time field of the neuron (**Fig. 1C**; see Materials and Methods for details). After being sorted by the time field, sequential activations of the neurons are evident as a diagonal dark band in the averaged spiking activity during the movie watching. This is especially strong for the visual (VIS) neurons and to a much less extent for the hippocampal CA1 neurons. The same analysis on the shuffled data, where neuronal spikes were temporally shuffled within each movie trial, results in much lighter diagonal bands (**Fig. 1D**, upper panels). Consistent with this observation, the peak time specificity scores derived from the original data are significantly higher than those of the shuffled data (**Fig. 1D**, lower panels). In addition, the diagonal bands from the original data are curved at the beginning and end of the movie, suggesting a disproportional representation of time, whereas those from the shuffled data are largely straight lines (**Fig. 1D**, upper panels). The distributions of the peak time specificity score were significantly different between the real and shuffled data, as measured by the Kolmogorov–Smirnov (KS) scores. The differences are much larger for the VIS neurons than for the CA1 neurons (**Fig. 1E**). Similar results were also obtained for the post-rest movie session and an extended group of mice (**Fig. S1C and S1D**). The neurons with a significant (*p* < 0.05) peak time specificity score were regarded as time-selective neurons. This is different from the conventionally defined time neurons (20, 22) since in this case their activity may have been responses to events in the movie stimulus rather than only reflect the passage of time. Both the CA1 and VIS time-selective neurons are reproducible across presentations of the same movie (**Fig. S1E and S1H**) with the CA1 time-selective neurons show more variability relatively (**Fig. S1E and S1G**).

### Movie-induced sequence of the CA1 time-selective neurons replays at rest

We then studied whether the firing sequence observed during the movie watching would replay at rest, similar to the place cell firing sequence during maze running (6, 7). We adapted a template matching method to detect the replay events. Briefly, the resting-state spiking data were divided into time segments according to troughs of the global mean spiking rate (vertical dotted lines in **Fig. 2A**) similar to the previous study (25), but the global mean signal was first low-pass (<5 Hz) filtered to generate fine-scale segments whose duration (556 ± 186 ms) roughly matched up with the known timescale of hippocampal replays. A delay profile was computed to describe the order of sequential activations of time-selective neurons within each segment, and then correlated (Spearman’s rank correlation) with the time-selective neuron firing sequence during the movie (**Fig. 2A** and **2B**; see Materials and Methods for details). The replay events were then detected as time segments showing significant (*p* < 0.01, horizontal dash lines in **Fig. 2A**) positive (forward) and negative (reverse) correlations (red and blue bars in **Fig. 2A**). The same procedure was repeated for randomized movie sequences (*N* = 200) to create a null distribution for the replay counts. In 8 out of 14 mice, the number of replay events of the CA1 time-selective neurons were significantly higher than what would be expected from the randomized controls. The above analysis was repeated for three control cases: the equal number of VIS time-selective neurons showing the strongest time-selective responses to the movie, the CA1 non-time-selective neurons that did not show significant time-selective responses, and the CA1 time-selective neurons derived from the shuffled data described above. Significant replay events were seen in none of these cases, including the VIS time-selective neurons that had a stronger time-specific responses in the movie sessions than the CA1 time-selective neurons. Consistent with the previous findings (6, 7, 28), independently detected SPW-R events (see Methods) peaked around the center of the replay events of the CA1 time-selective neurons (**Fig. 2E**).

**Figure 2.**
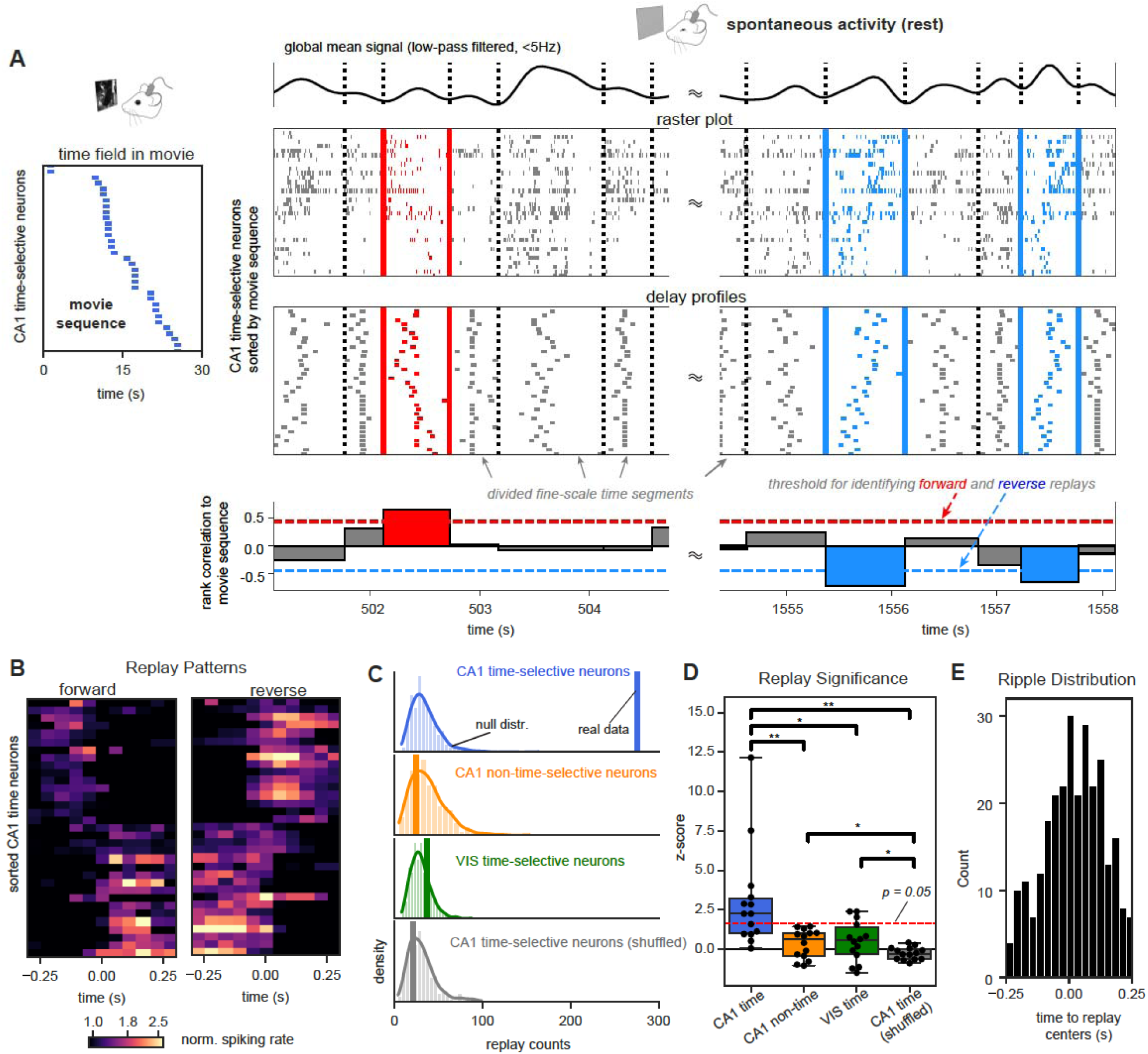
Movie-induced sequence of hippocampal time-selective neurons replayed at rest. **(A)** Detection of replays in a representative mouse. Spiking data was divided into fine-scale time segments according to troughs of the filtered (<5 Hz) global mean signal (top). The segment boundaries were marked by dotted lines. A delay profile (the 3rd row) was computed to describe the relative timing of the time-selective neurons’ spiking activity (the 2 rd row) within each time segment. A template of movie-induced sequence (left) was constructed based on the time fields of the CA1 time-selective neurons in the movie. The bar plot (bottom) shows the Spearman’s rank correlations between the movie sequence and the delay profiles of the fine-scale segments. The forward (red) and reverse (blue) replays were detected as the time segments showing significant (p < 0.01) positive and negative correlations, respectively. **(B)** The averaged pattern of the forward (left) and reverse (right) replays from the representative mouse. They were obtained by aligning and averaging the detected replay segments. **(C)** The number of detected replay events was compared against a null distribution built with repeating the same analysis on randomized movie sequences. The same result was derived for the CA1 time-selective neurons, CA1 non-time-selective neurons, VIS-time-selective neurons, and CA1-time-selective neurons identified from the shuffled data (from top to bottom). **(D**) The box plot of z-scores quantifying the difference between the real counts of replay events and the null distributions from all 14 mice. **(E)** The distribution of hippocampal ripples counts relative to the detected replay events of the CA1 time-selective neurons from all 14 mice.

### Hippocampal replays co-occur with brain-wide spiking cascades

We further investigated the potential link between the replay events and previously reported brain-wide cascades of neuronal firing (25). The slow spiking cascades can be clearly seen in the resting-state recordings after sorting all recorded neurons from various brain regions according to their principal delay profile (**Fig. 3A**), i.e., the first principal component of delay profiles of coarse-scale time segments (see (25) for more details). This coarse-scale principal delay profile represents the direction of sequential activations of the spiking cascade. The cascade started with slow and sequential entrainments of the negative-delay neurons (top, blue-symbolled neurons in **Fig. 3**) at the early phase and then reached a tipping point featuring the rapid transitioning to the activation of the positive-delay neurons (bottom, red-symbolled neurons in **Fig. 3**), which were then slowly and sequentially disengaged in ∼1-3 seconds (**Fig. 3A** and **3B**). The cascade involved ∼70% of all recorded neurons from various brain regions (25), and the region-specific mean spiking activity showed significant modulations at the cascade in every recorded brain region (**Fig. 3C**). Tracking the occurrence of the CA1 replays along with the spiking cascade revealed an interesting pattern: the reverse replays of movie sequence in the CA1 time-selective neurons are much more likely to occur around the fast transitioning (yellow arrows) of the spiking cascades (**Fig. 3A**). This observation is consistent with the distribution of the reverse replays over the cascade cycle (**Fig. 3D** and **3E**). The forward replays displayed an opposite modulation and were less likely to appear around this transitioning point (time zero in **Fig. 3D** and **3E**). In comparison, the replays detected for the three control groups of neurons, including the VIS time-selective neurons, did not show significant modulations across the spiking cascade cycle, particularly at the transitioning point (**Fig. S4E**).

**Figure 3.**
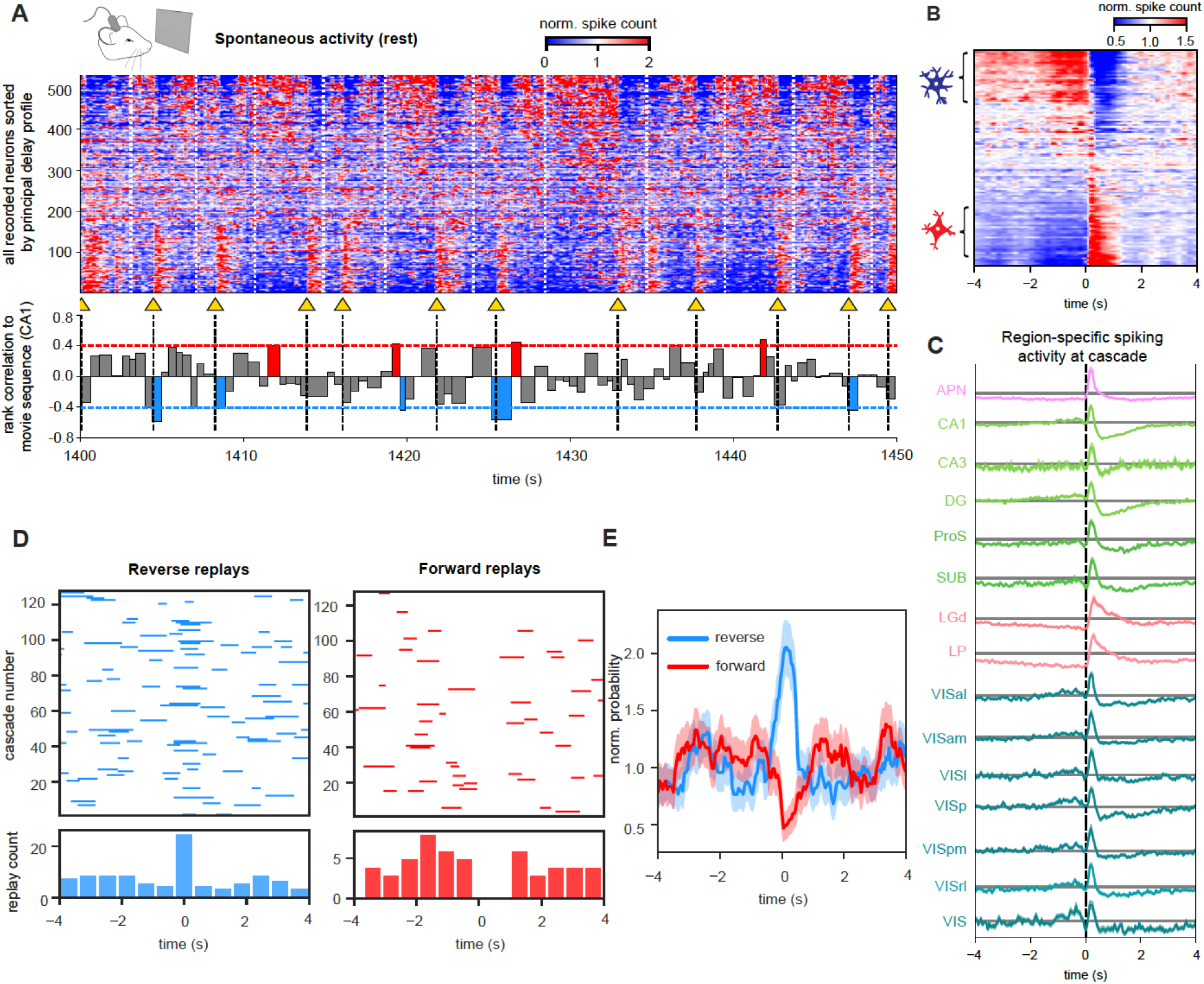
Hippocampal replay events temporally locked to spiking cascades across the forebrain. (**A**) A 50-s example of spiking data during the resting state in a representative mouse with all recorded forebrain neurons being sorted by the principal delay profile (top). Boundaries of the coarse-scale time segments and spiking cascade (dashed white lines) were delineated by the troughs of the coarsely filtered (<0.5 Hz) global mean signal. The bar plot (bottom) shows the rank correlations between the movie sequence and the delay profile of the CA1 time-selective neurons. The forward and reverse replays were colored by red and blue respectively, and the dotted horizontal lines represent the thresholds for detecting the replay events. Yellow arrows and black dotted lines mark the fast-transitioning points from the ‘negative-delay neurons to the positive-delay neurons. (**B**) The averaged pattern of the spiking cascade from the representative mouse. (**C**) Averaged spiking dynamics of different brain regions at the slow spiking cascade. Only 15 brain regions with > 100 neurons were shown. (**D**) The detected reverse (top left) and forward (top right) replays were distributed over the cycle of the spiking cascades from a representative mouse. Each row corresponds to a spiking cascade and short horizontal lines represent the detected replays. The length of each line equals to the duration of the replay. Their distributions were summarized in the histograms (bottom). (**E**) The normalized probability of forward (red) and reverse (blue) replays across the cascade cycle with the data from all the 14 mice. The shaded region denotes area within 1 SEM (N = 1787).

### Distinct micro-cascades mark forward and reverse replay events

Our pilot investigations revealed similar cascade dynamics of shorter sub-second timescale, which we here term “micro-cascades”, and related them to both the occurrence and structure of spontaneous replay events. The structure of such micro-cascades can be seen in Fig 4A-D. Briefly, these finer-scale events featured a similar sequential transition from the negative-delay neurons to the positive-delay neurons as the coarse-scale cascade, but the positive-delay neurons were only briefly activated for <100ms. Importantly, they were often associated with single SPW-R events (**Fig. 4B**, red arrows). To better understand the fine-scale dynamics, we correlated the delay profiles of the fine-scale time segments with the coarse-scale principal delay profile. The resulting sequential scores (i.e., normalized correlations) were significantly (KS test; *p* = 0) stronger than randomized controls (**Fig. 4C**). Unlike the sequential scores of the coarse-scale segments that mostly showed large positive values (**Fig. S4A**), the fine-scale segments have both large sequential scores of negative and positive values (**Fig. 4A**). In addition, the principal delay profile derived directly from the fine-scale segments is highly similar to the coarse-scale principal delay profile (**Fig. S6B**), suggesting that both slow (seconds) and fast (hundreds of milliseconds) cascade dynamics feature sequential activations along a similar direction.

**Figure 4.**
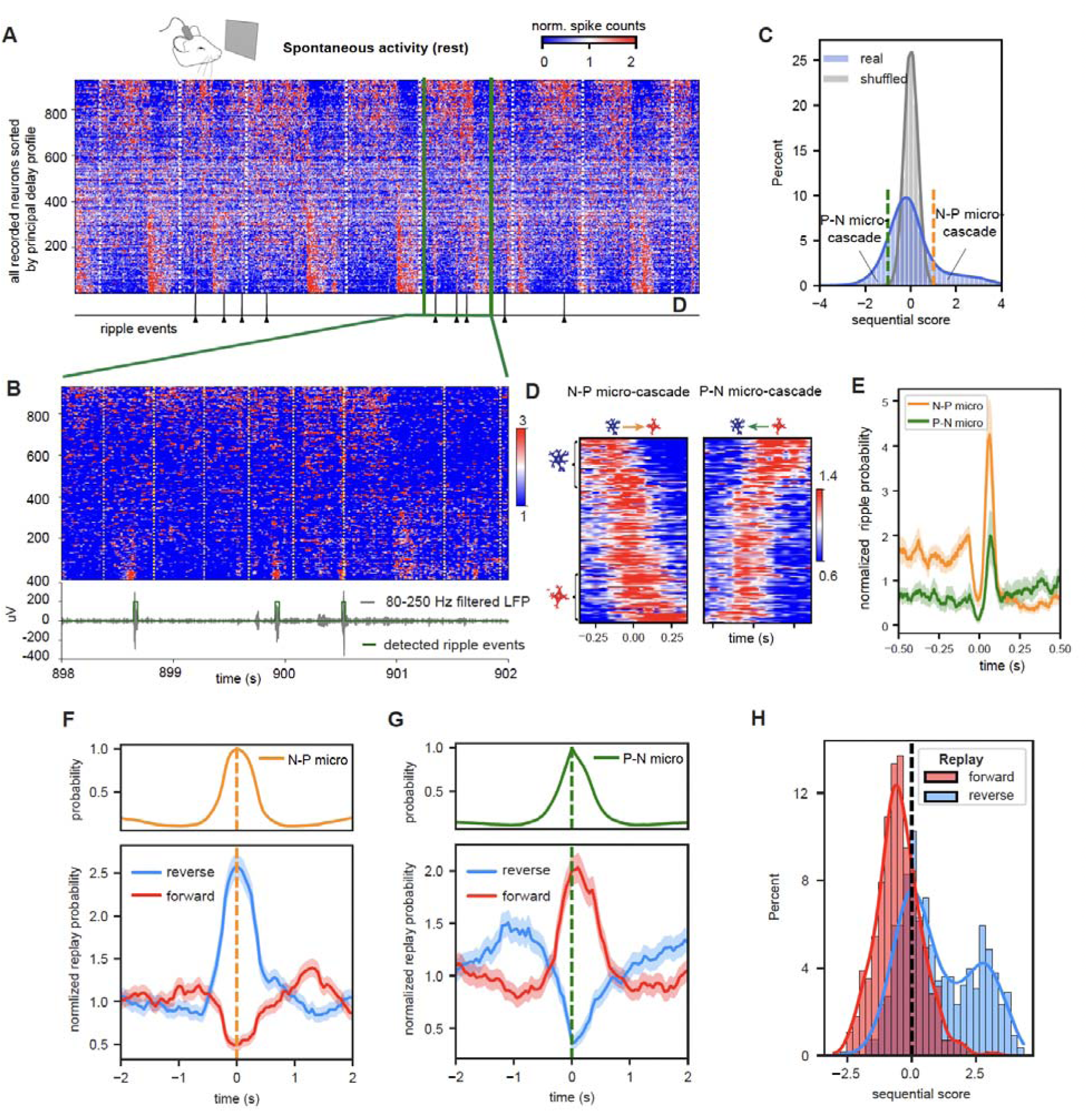
Distinct micro-cascades are associated with the forward and backward hippocampal replay events. (**A**) An example of resting-state spiking data with a finer (20ms) temporal resolution. The bottom trace shows the identified SPW-R events. (**B**) A 4-sec segment in (A) was amplified horizontally. The spiking data was divided into fine-scale segments based on the troughs of the finely filtered (<5 Hz) global mean signal. The period with apparently sustained activations of the negative-delay neurons is punctuated by very brief (< 100ms) activations of positive-delay neurons, which are often associated with a single SPW-R. (**C**) The distribution of sequential scores of all the fine-scale segments. The sequential score is the normalized correlation between the delay profile of the fine-scale segments and the coarse-scale principal delay profile. The fine-scale segments with significant (p < 0.001) positive (right to the orange dash line) and negative (left to the green dash line) sequential score were defined as the negative-to-positive (N-P) and positive-to-negative (P-N) micro-cascades. (**D**) The averaged patterns of the N-P (left) and P-N (right) micro-cascades from a representative mouse, aligned and averaged according to the global spiking peaks in the identified micro-cascades. (**E**) The normalized probability of SPW-R across the cycle of the N-P and P-N micro-cascades. They were aligned and averaged according to the brief peaks of positive-delay neuron activations in the micro-cascades. (**F, G**) The normalized probabilities for the forward and reverse replays across the cycle of the N-P and P-N micro-cascades. (**H**) Sequential score distributions for the fine-scale segments associated with the forward and reverse hippocampal replays.

We then extracted the fine-scale segments with significant (*p* < 0.001) negative and positive sequential scores and called them the P-N (positive-delay neurons to negative-delay neurons) and N-P micro-cascades respectively. Their averaged patterns clearly showed sequential activations along and opposite to the principal delay profile direction (**Fig. 4D**). The brief positive-delay neuron activation at these micro-cascades was tightly coupled by a sharp increase in the SPW-R probability (**Fig. 4E**). Most importantly, the reverse and forward replays co-occurred with the N-P and P-N micro-cascades respectively (**Fig. 4F** and **4G**). At the same time, the sequential scores of the reverse and forward replay segments are biasedly distributed towards the negative and positive values respectively (**Fig. 4H**). These results remained similar with removing the micro-cascades, mostly the N-P type, at the fast transitioning point of the slow spiking cascades (**Fig. S7**).

## Discussion

Here we examined the activity of a large population of neurons from throughout the brain during hippocampal replay following passive movie viewing in rodents. We found that both forward and reverse hippocampal replay were embedded within brain-wide cascades of sequential neuronal activation involving many forebrain structures. Within the replay activity, the reverse hippocampal replay events were most directly correlated with the peaks of these large-scale cascades. At a finer timescale, both forward and reverse replay events matched unique brain-wide cascade patterns.

The embedding of hippocampal replays in the highly structured, resting-state global dynamics supports recent theory about the self-organized nature of the hippocampal neural sequences (12, 15, 20). The replays of movie-related hippocampal sequence observed here are similar to what has been repeatedly reported for maze-running-related place-cell sequences (21). Interestingly, the place-cell sequences were also found to “pre-play” before the maze running. While such pre-plays had once been explained as the internal dynamics for action planning (21), this planning interpretation may not explain the pre-plays occurring even before animals see the maze track (17, 19). The pre-play finding is however consistent with another line of research into hippocampal time cells (14, 20, 29) since both suggested the self-generated nature of hippocampal sequences. It was found that hippocampal neuronal sequences can be robustly formed with animals running on a wheel without apparent changing of environmental or body-derived inputs, suggesting that they actually represent self-generated dynamics for time-encoding (20).

The existence of apparent time cells has led to the idea that the sequential firing in the hippocampus during a temporally structured event may be internally generated rather than driven by a sequence of external stimuli(12, 20). The new theory would reconcile the “pre-play” and “re-play” findings if self-generated sequential dynamics generally follow a pre-existing temporal order. Here we showed that the movie evoked the time-selective responses, and thus the temporal activation sequences, of both the hippocampal and visual neurons. The sequence of the hippocampal time-selective neurons, but not the visual neurons with stronger time-selective responses, was found to replay during the rest period after the movie watching. The difference might be due to the fact that the time sequences in hippocampus result from the firing order imposed by its neural substrate, while the order observed in visual cortex is imposed by time-specific movie features.

Importantly, the replays of the hippocampal sequences were embedded in pre-existing, self-organized global brain dynamics, consisting of coarse- and fine-scale spiking cascades (25). These resting-state activity cascades featured sequential activations of the whole-brain neuronal populations along a specific direction. This temporal direction governed the sequential activations of different timescales and across different populations, including the hippocampal sequence during the movie watching (30 sec), the coarse spiking cascades (5-15 sec), the micro-cascades and the replays of the movie sequence (∼hundreds of milliseconds). Thus, it may represent a general direction of sequential activity in the brain.

Increasing evidence suggested that the hippocampal ripples are coordinated with brain-wide neural dynamics (25, 30–32). The present study extended these findings by showing that the hippocampal replays are embedded in the global cascade dynamics of sequential activation. This arrangement could have certain advantages at least theoretically.

First, the global dynamics may open a critical time window for the hippocampo-cortical interactions that are essential for memory consolidation. The spiking cascades involved ∼70% of brain neurons in various cortical and subcortical areas. Particularly around the rapid transitioning point, most of the recorded neurons, including the hippocampal and cortical neurons, fired within a very brief (hundreds of milliseconds) time window, and created an opportunity for information transfer between the hippocampus and the cortex. The hippocampo-cortical interplay has been observed previously as slow (∼10 sec) co-modulations of the cortical delta-band power and the hippocampal ripples (33, 34). The ripples were also found to trigger widespread cortical fMRI responses of the seconds timescale (35). These hippocampo-cortical interactions may represent the same brain process as the spiking cascade, which was coupled to slow modulations of both the cortical delta power and hippocampal ripples (25).

Second, the embedding of the hippocampal replays in the global dynamics could be an efficient way of consolidating the learning and memory. Different daily-life experiences can be encoded in neuronal sequences of different subgroups of hippocampal neurons (15, 21, 24). The spiking cascades that entrain most brain neurons would then be able to replay them all at once through a global sequential activation following the pre-existing principal direction.

Lastly, the global spiking cascades may provide the driving forces for the hippocampal replays. The importance of the hippocampal replays makes their occurrence unlikely to rely completely on random fluctuations of spontaneous brain activity. In the absence of external perturbations during rest and sleep, the self-generated dynamics could be critical for driving these events in a controllable way. The highly organized spiking cascades would serve this purpose by driving the replay events and warranting their re-occurrences. Nevertheless, it remains unclear what in turn drives the spiking cascades. Modulatory influences from the various neurotransmitter systems, including the cholinergic system (36–38), are among the possibilities. The resting-state global brain activity measured by fMRI and electrocorticography has been linked to subcortical arousal-regulating areas (39), particularly the major locations of the cholinergic neuron (40, 41). In fact, the deactivation of the basal forebrain cholinergic regions effectively suppressed the resting-state global activity. The spiking cascades, which are shown as the global brain activity of single neuron level, were phase coupled to slow pupil dilations (25), which have previously been shown to be linked to the activation of cholinergic neurons (42). This explanation would be consistent with the known role of the cholinergic projections in the generation of the hippocampal ripples (43–45).

## Acknowledgements

This work was supported by the Brain Initiative award (1RF1MH123247-01), the NIH R01 award (1R01NS113889-01A1), and the Intramural Research Program of the National Institute of Mental Health (ZIA-MH002838). We also thank Dr. Feng Han for proof reading the paper and assisting some figure illustrations.

## Data and materials availability

We used the Neuropixels Visual Coding dataset from the Allen Institute (26, 27). All the multimodal data are available at https://portal.brain-map.org/explore/circuits/visual-coding-neuropixels. The Python code that produced the major results of this paper will be available at https://github.com/psu-mcnl/Neural-Seq.

