## supplementary figures and methods for "Self-generated brain-wide spiking cascades govern replay dynamics in the hippocampus"

**Affiliations:**

**Author Contributions:**

**Y.Y.&X.L.** contributed to the conception, design of the work, and data analysis;
**X.L.** also devoted the efforts to the supervision, project administration and funding acquisition;
**Y.Y., D.A.L., J.H.D. & X.L.** contributed to data visualization, and writing the paper.

### Supplementary Figures

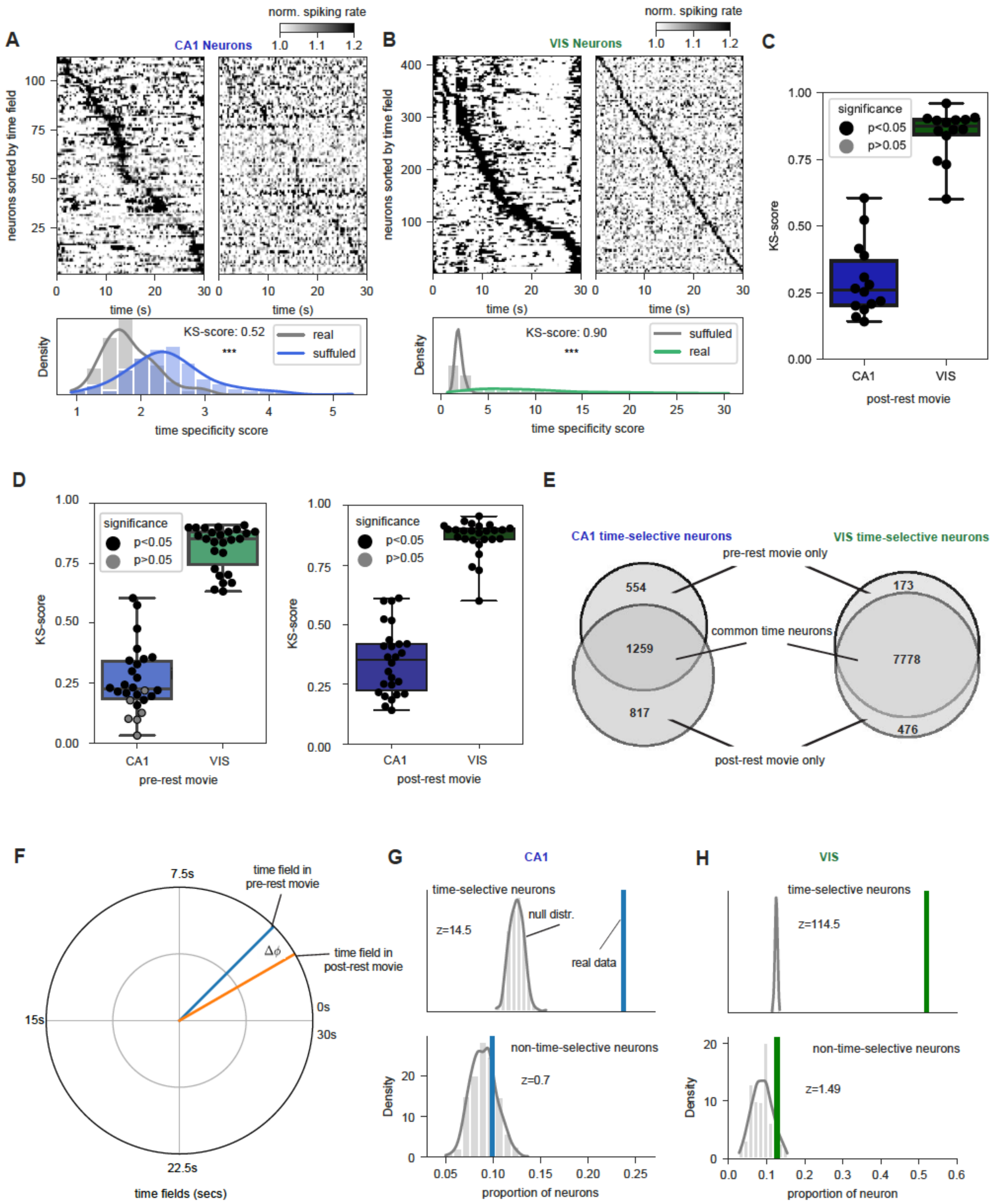


***Figure S1. Time-specific responses of the hippocampal and visual neurons.*** *(A) The averaged activation pattern of neurons from CA1 regions over 30 trails in post-rest movie sorted*

*according to neuron's time field for an example mouse (top) where the results from original data (left) and shuffled control (right) were shown. Distribution of the peak time specificity scores were compared between the real and shuffled data (bottom). The similar analysis was also performed for VIS neurons in post-rest movie (B). (C) The box plot of Kolmogorov-Smirnov (KS) scores summarized for both CA1 and VIS neurons across all 14 mice in post-rest movies. (D) Similar results were also summarized for all 26 mice in pre-rest movies (left) and post-rest movies (right). (E) The Venn diagrams of the number of time-selective neurons under three conditions: only pre-rest movies, both pre- and post-rest movies, and only post-rest movies. It was summarized for the CA1 regions (left) and VIS regions (right), respectively. (F) Illustration of time fields consistency quantification between pre- and post-rest movies. The time fields were projected to polar coordinates and the phase difference quantifies the variation between the time fields pair for a neuron. (G) The proportion of a neuron group showing the consistent time fields between pre- and post-rest movies was compared against the null distribution built by repeating the same analysis on the same group of neurons with randomly shuffled time fields. We derived results for CA1 common time-selective neurons (top) and CA1 common non-time-selective neurons (bottom), which are defined respectively as the time- or non-time-selective neurons common to both pre-rest and post-rest movies. (H) Similar results were obtained for the visual regions.*

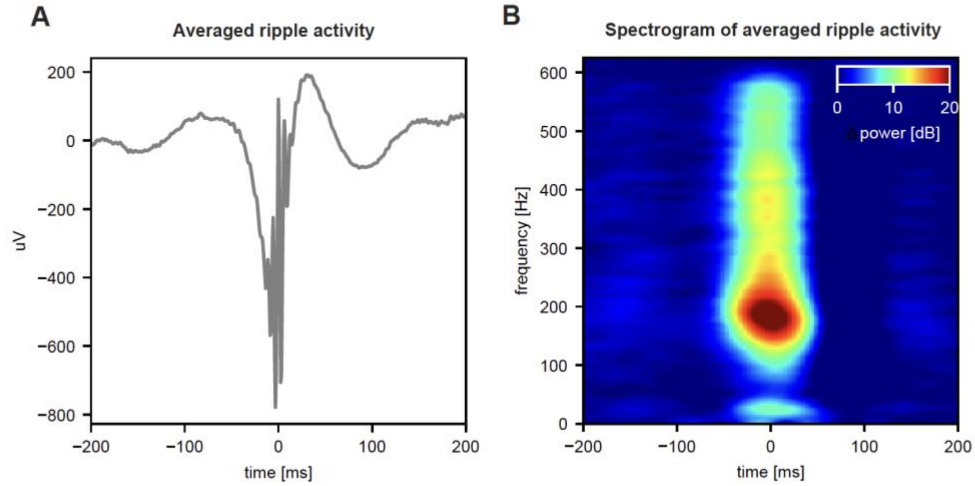


***Figure S2. Hippocampal ripple detection.*** *(A) The averaged LFP signals across all detected ripple events for a representative mouse. (B) The spectrogram of LFP was computed and averaged across all detected ripple events for a representative mouse.*

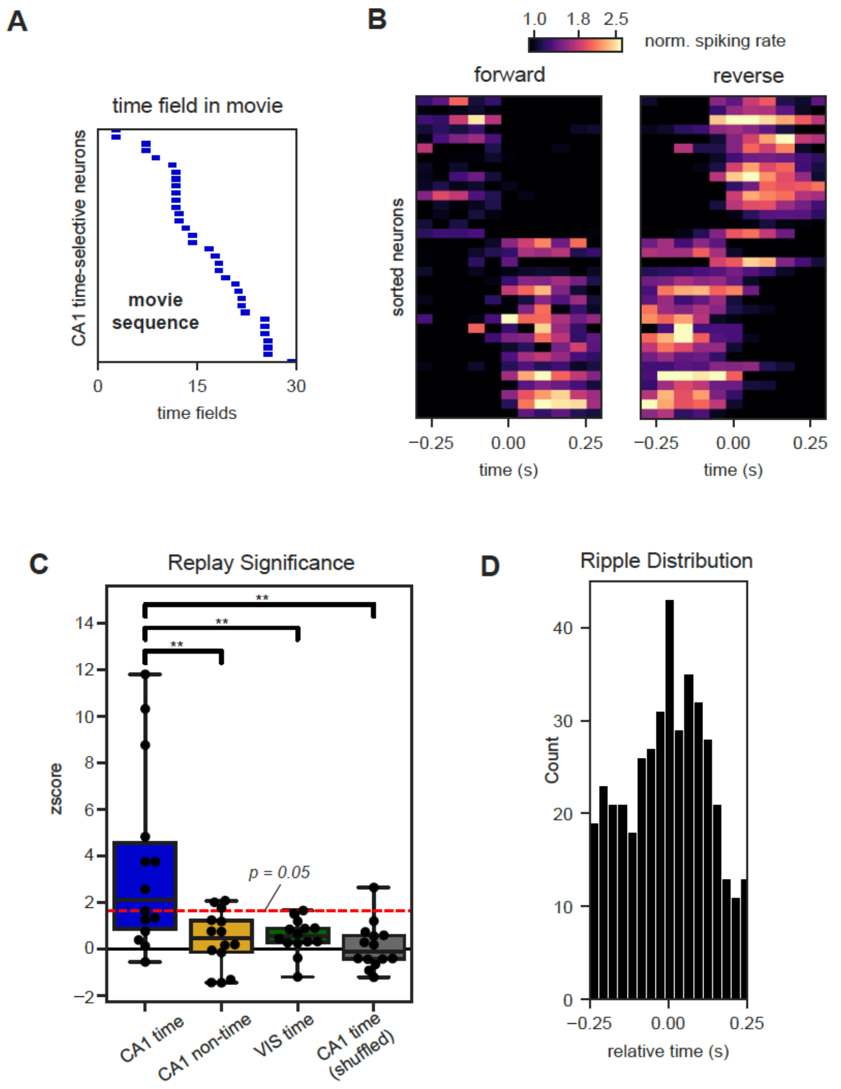


***Figure S3. Post-rest movie sequence replayed at rest.*** *(A) The template of post-rest movie sequence was constructed based on the CA1 time-selective neurons. (B) The averaged pattern of the forward (left) and reverse (right) replays of the post-rest movie sequence in the representative mouse. They were obtained by aligning and averaging the detected replay segments. (C) The box plot of z score quantifies the difference between the null distribution and the real observation in terms of the replay count of the post-rest movie sequence from all 14 mice. (D) The distribution of hippocampal ripples counts relative to the detected replay events of the post-rest movie sequence from all 14 mice.*

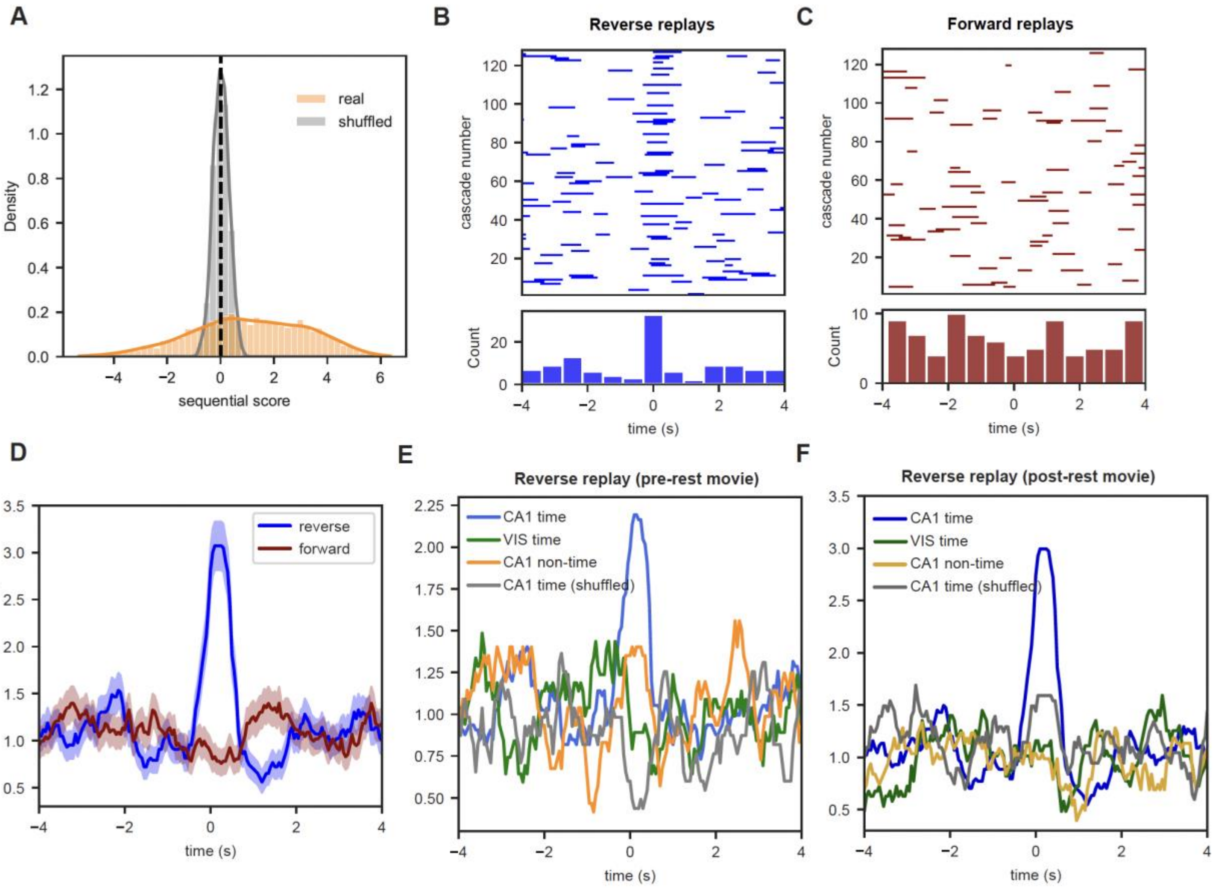


***Figure S4. Reverse replays of the movie sequences occur preferentially at the fast-transitioning point of the spiking cascade.*** *(A) The distribution of sequential scores for all coarse-scale time segments. They are summarized for both real data and randomly shuffled data. Reverse (B) and forward (C) replays of the post-rest movie sequence were distributed over the cycle of the spiking cascades from a representative mouse (top). The histograms (bottom) summarized the distribution of the replay events for this representative mouse. (D) The normalized probability of forward (dark red) and reverse (dark blue) replays across the cascade cycle with the data from all 14 mice. (E) The normalized probability of reverse replays of the pre-rest movie sequence at cascade for the 4 different groups of neurons: CA1 time-selective neurons, CA1 non-time-selective neurons, VIS-time-selective neurons, and CA1-time-selective neurons identified from the shuffled data. (F) Similar results were summarized for the reverse replays of the post-rest movie sequence.*

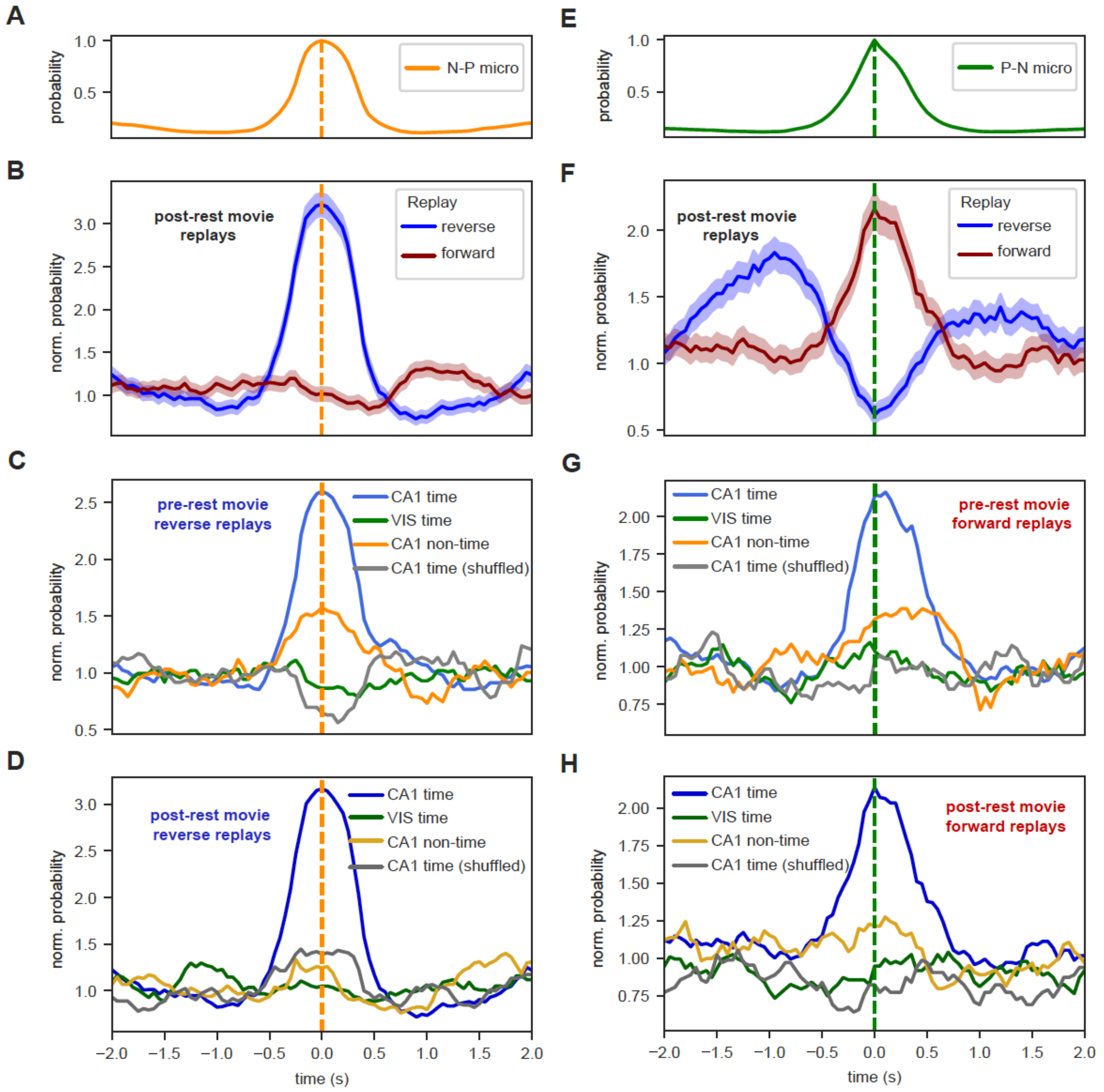


***Figure S5. Distinct micro-cascades are associated with different types of hippocampal replays.*** *(A) The N-P micro-cascade probability, (B) the normalized probability of reverse and forward replays of the post-rest movie sequence (i.e., for CA1 time-selective neurons) at N-P micro-cascade, and the normalized probability of reverse replays of the pre-rest (C) and post-rest (D) movie sequences for the 4 different neuron groups: CA1 time-selective neurons, CA1 non-time-selective neurons, VIS-time-selective neurons, and CA1-time-selective neurons identified from the shuffled data. (E-H) Similar results were obtained for the P-N micro-cascade probability.*

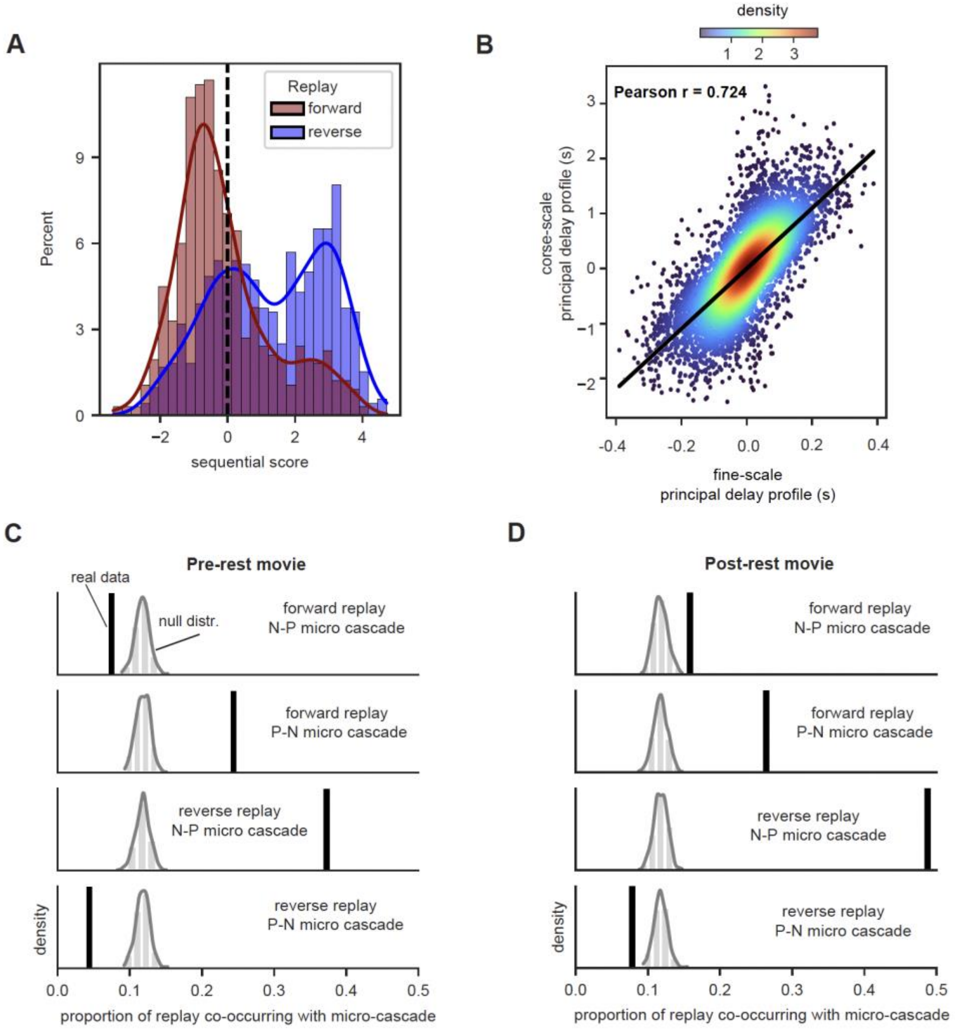


***Figure S6. Hippocampal replays are associated with micro-cascades.*** *(A) Sequential score distributions for the fine-scale segments associated with the forward and reverse hippocampal replays of the post-rest movie sequence. (B) The scatter plots showing the linear relationship between the principal delay profile of the coarse-scale time segments and that of the fine-scale time segments. (C) The proportion of the hippocampal replays co-occurred with the detected micro-cascades for the pre-rest movie sequence (black vertical lines) and its randomly shuffled versions (500 repeats, gray distributions). The result was derived for four conditions corresponding to four different combinations of types of replay and micro-cascades, i.e., forward replay vs. N-P micro cascades. For all conditions, there was a significant difference between real and control, with* $p < 10^{-5}$*. (D) Similar results were summarized for the post-rest movie sequence and the difference is significant for all conditions (*$p < 10^{-5}$*).*

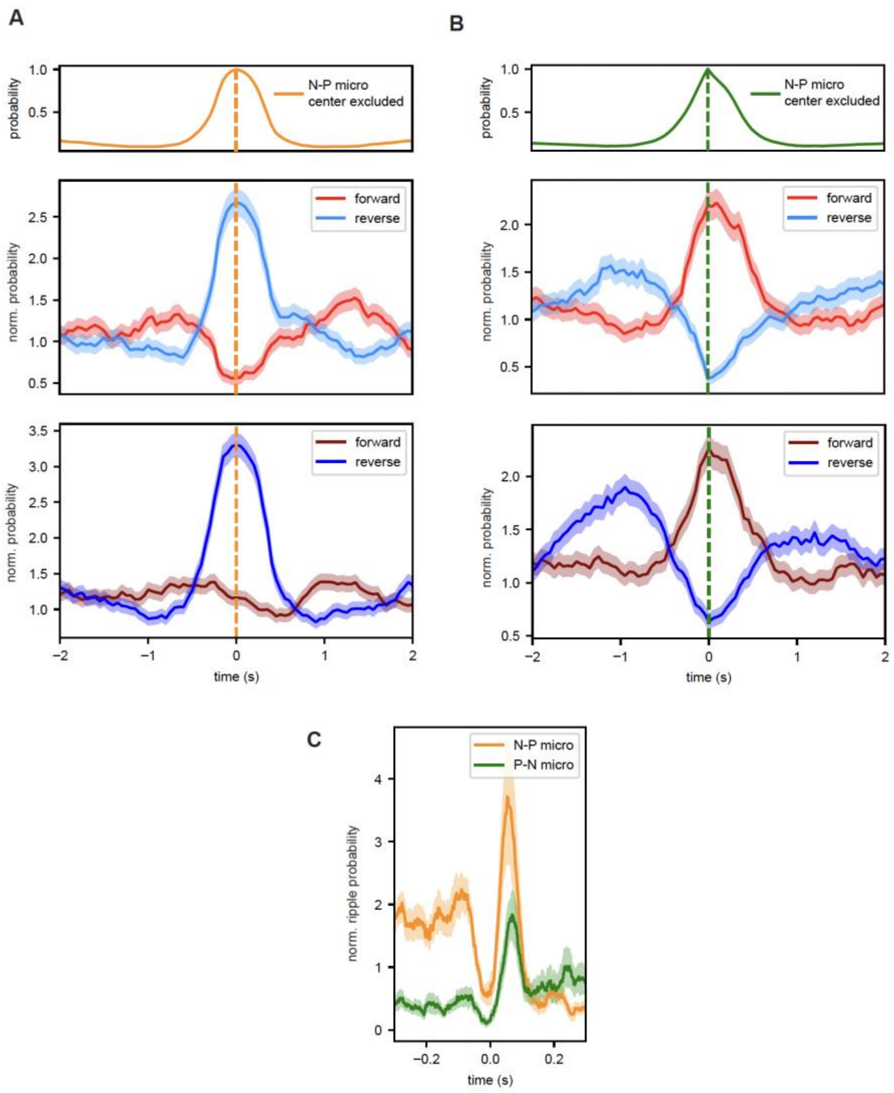


***Figure S7. Micro-cascades off the fast-transitioning point of the slow cascades.*** *The normalized probability of the forward and reverse replays across the cycle of a subgroup of N-P micro-cascades (A) and P-N micro-cascades (B) that were not overlapped with those detected at the fast-transitioning point of the slow cascade (i.e., those shown in **Figure 3**). The results were summarized for the pre-rest (middle) and post-rest movie (bottom) movie sequences, respectively. (C) The normalized probability of SPW-R across the cycle of this subgroup of the N-P and P-N micro-cascades.*

### Materials and Methods

#### Neuropixels data

The present study was performed on the Neuropixels Visual Coding dataset of the Allen Institute (1, 2). The dataset contains neural spiking data recorded from mice with the advanced Neuropixels technique and behavioral measurements including running speed and eye tracking. The probes were capable of recording neural activity from cortical areas to subcortical areas, including hippocampal formation (CA1, CA3 and dentate gyrus).

The dataset contains two stimulus sets, the "Brain Observatory 1.1" and the "Functional Connectivity." We focused on the "Functional Connectivity" stimulus set which contains a 30-minute spontaneous session and several standardized sets of visual stimuli including two natural movie sessions. The two natural movie sessions (pre- and post-rest movie sessions) were presented before and after the spontaneous session. For each natural movie session, a 30-second clip from the movie *Touch of Evil* was passively presented to mice repeatedly for 30 times. There are 26 mice from the "Functional Connectivity" stimulus set, including three transgenic lines (Sst-164 IRES-Cre, Vip-IRES-Cre, and Pvalb-IRES-Cre) and wild type. The time-selective neuron analysis was performed for all mice. We mostly focused on 14 mice with sufficient immobile periods in spontaneous sessions (>10 min) and eye-tracking data for replay analysis.

#### Neural spikes

The neural spikes data were sorted with a pipeline that implemented Kilosort2 (3) for spike sorting and the details of the pipeline can be found in the original papers (1, 2).

To compute neural spike rate, we evenly divided time into time bins and counted neural spikes within each time bin. We used different sets of time bins to detect different time-scale events. For fast-scale events such as replay and micro-cascade, the size of each time bin is 50 ms. For slow-scale events such as the slow cascade, the size of each time bin was set to 200 ms, following the setting in a previous paper (4). Then, the spiking rate of each neuron was normalized by its temporal mean spike rate. The global mean spiking rate was computed by averaging the normalized spiking rate over all recorded neurons.

#### Time-selective neurons identification

To investigate the existence of time-selective neurons in the movie sessions, we developed a method to measure the time-specific response for each neuron. First, the 30-second movie was evenly segmented into $N_{TB}$ time bins with $N_{TB} = 60$. Then, for each time bin $j$ and for each neuron $i \in U$, we formed a response vector $r_{ij}$ by grouping the mean spiking rate of this neuron at this time bin for all 30 movie repeats. We also formed a baseline vector $b_{ij}$ by grouping the mean spiking activity rate of this neuron at all other time bins for all 30 repeats. The two-sample t-test was then used to quantify is there a significant difference between the response vector $r_{ij}$ and baseline vector $b_{ij}$, and the resulting t-score was termed time-specificity score $s_{ij}$:

$$s_{ij} = \text{2-sample t-test}\left(r_{ij}, b_{ij}\right)$$

where $r_{ij}, b_{ij} \in \mathbb{R}^{30}$ and $s_{ij} \in \mathbb{R}$.

As a result, each neuron has a time-specific score at a specific time bin. We then defined a time field $TF_i^*$ of each neuron as the time bin it achieved the maximum time specificity score $s_i^*$.

$$TF_i^* = \text{argmax}_{1 \le j \le N_{TB}} s_{ij}$$

$$s_i^* = \max_{1 \le j \le N_{TB}} s_{ij}$$

A neuron was regarded as a time-selective neuron with a significant time-specific response if its maximum time-specific score $s_i^*$ was larger than a pre-set threshold $\gamma = 2$, which was approximately corresponding to a significance level of $p = 0.05$ (two-tails 2-sample t-test, DF=58). The set of time-selective neurons $U_T$ from a mouse was defined as

$$U_T = \{i\colon \mid s_i^* > \gamma, \text{ for } i \in U\}$$

For the hippocampal CA1 region, a set of CA1 time-selective neurons was defined as $U_{\text{CA1-T}} = U_T \cap U_{\text{CA1}}$. Accordingly, time-selective neurons in the visual region were defined as $U_{\text{VIS-T}} = U_T \cap U_{\text{VIS}}$. Based on the neurons' preferred time field, we constructed a movie sequence $I_{\text{mov}}$ which defines the firing order of neurons in the movie. For example, we defined the movie sequence $I_{\text{mov-T}}$ as:

$$\begin{aligned} I_{\text{mov-T}} &= \text{argsort}_{i \in U_{\text{T}}} TF_i^* \\ &= \{i \dots j \dots k \in U_{\text{T}} \mid TF_i^* \le \cdots \le TF_j^* \le \cdots \le TF_k^*\} \end{aligned}$$

Similarly, for each subset of time-selective neurons, we defined their movie sequence accordingly, $I_{\text{mov-CA1-T}}$ for CA1 time-selective neurons and $I_{\text{mov-VIS-T}}$ for VIS time-selective neurons.

To build a control dataset, for each movie session we randomly shuffled the neural activity of each neuron within each repeat of the movie over the $N_{TB}$ time bins. By applying the same procedure of time-selective neuron identification to the shuffled data, we obtained corresponding time-selective neurons $U_{T^s}$, CA1-time-selective neurons $U_{\text{CA1-T}^s}$, VIS-time-selective neurons $U_{\text{VIS-T}^s}$, movie sequences $I_{\text{mov-T}^s}$, $I_{\text{mov-CA1-T}^s}$ and $I_{\text{mov-VIS-T}^s}$. We compared the distribution of the time-specificity scores between the original and shuffled data for both CA1 and VIS neurons. Their difference would quantify whether the ensemble of neurons contains time-selective neurons. We quantified the difference using Kolmogorov-Smirnov (KS) test that is a nonparametric test of the equality of two data distributions.

$$d_{\text{CA1}} = \text{KS-score} = \text{KS-test}\left(\{s_i^*\}_{i \in U_{\text{CA1}}}, \{s_i^*\}_{i \in U_{\text{CA1}}}^s\right)$$

$$d_{\text{VIS}} = \text{KS-score} = \text{KS-test}\left(\{s_i^*\}_{i \in U_{\text{VIS}}}, \{s_i^*\}_{i \in U_{\text{VIS}}}^s\right)$$

where $\{s_i^*\}^s$ represents the set of time-specificity scores derived for shuffled movie response.

For another control, we also found another set of non-time-selective neurons in the CA1 region, defined as $U_{\text{CA1-nT}} = U_{CA1} \backslash U_{\text{CA1-T}}$.

**Time field consistency of the time-selective neurons**

We used two movie sessions, i.e., the pre- and post-rest movie sessions in our analysis. We identified time-selective neurons for each movie session independently and obtained two versions of time-selective neurons and corresponding time fields

$$U_T, \{TF_i^*\}_{i \in U_T} \quad \text{for pre-rest movie,}$$
$$U'_T, \{TF_i^{*'}\}_{i \in U'_T} \quad \text{for post-rest movie.}$$

The movie was repeatedly played for 30 times in each session. To understand how the set of time-selective neurons and their time field changes across the two sessions, we re-represented the time field as a polar angle. For $i \in U_T$,

$$\phi_i = 2\pi \frac{TF_i^*}{N_{TB}}$$

Similarly, for $i \in U'_T$, we applied the same transformation

$$\phi'_i = 2\pi \frac{TF_i^{*'}}{N_{TB}}$$

The time field change of a neuron between the two movie sessions was then measured by the phase difference:

$$\Delta\phi_i = |\phi_i - \phi'_i|$$

We considered the time field of a neuron is consistent when the phase difference smaller than a pre-set threshold $\phi_{thr}$ that is equal to 3 times phase resolution or $3\frac{2\pi}{N_{TB}}$.

We then found common sets of the time-selective neurons and non-time-selective neurons in the two movie sessions for both CA1 and VIS regions:

$$U_{CA1-T-com} = U'_{CA1-T} \cap U_{CA1-T}$$
$$U_{VIS-T-com} = U'_{VIS-T} \cap U_{VIS-T}$$
$$U_{CA1-nT-com} = U'_{CA1-nT} \cap U_{CA1-nT}$$
$$U_{VIS-nT-com} = U'_{VIS-nT-T} \cap U_{VIS-nT-T}$$

For each of these neuron sets, we evaluated the proportion of neurons showing the consistent time fields. Take the set $U_{CA1-T-com}$ as an example, the proportion can be derived by

$$p_{CA1-T-com} = \frac{|\{i \in U_{CA1-T-com} \mid \Delta\phi_i < \phi_{thr}\}|}{|U_{CA1-T-com}|}$$

To analyze the significance, we performed permutation test for the four defined neuron sets. We randomly permuted the time fields obtained from both movie sessions and computed the resulting proportion of neurons showing a consistent time field. The process was repeated for 500 times to construct the null distribution. We then quantified the deviation of real proportion of consistent neurons from the null distribution by compute the z-score and associated p-value. The results suggested that the time field of time-selective neurons is significantly consistent compared to random control in both CA1 and VIS regions (**Fig. S1G and 1H**).

**Resting stationary state**

Following the definition of stationary state in the previous study (4), we identified the time periods when mice remained stationary based on the recorded running speed data. First, we applied a low-pass filter (3rd-order Butterworth filter with critical frequency at 0.05 Hz) to the running speed data to remove high frequency noise. We assumed that negative running speed resulted from noise. Hence, we found all the negative speed values, took the absolute value, and set its 0.05 percentile as a threshold. Any time point with running speed larger than this threshold was considered non-stationary. There were short and brief stationary periods intermixed with non-stationary periods. We only preserved stationary periods with a duration longer than 50 seconds to focus on more sustained resting state.

**Delay-profile decomposition**

A delay-profile decomposition method, first proposed in fMRI data (5), has been applied to neural spiking data to map the relative temporal relationship within a group of neurons (4). Briefly, we segmented the spiking rate data based on troughs of the filtered global mean spiking rate. We conducted the delay profile decomposition method at both fine and coarse time scales with setting the low-pass filter of the global mean spiking rate to 5 Hz and 0.5 Hz respectively. Each time segment then included a local peak of the global mean signal and thus may represent a neuronal event. For each neuron $i$ in time segment $j$, we computed its activation time $\widehat{t_{\mathrm{ij}}}$ as the temporal centroid of the firing rate within the time segment:

$$\widehat{t_{\mathrm{ij}}} = \frac{\sum_{\tau \in T_j} \tau \mathrm{f_i}(\tau)}{\sum_{\tau \in T_j} \mathrm{f_i}(\tau)},$$

where $T_j$ contains relative time points for the time segment $j$ and $f_i(\tau)$ is the firing rate of neuron $i$ at time $\tau$. Then, a delay profile $d_j$, which encodes relative temporal ordering of neurons within the time segment, was constructed for each time segment $j$ by grouping the activation time for all the neurons:

$$d_j = \begin{pmatrix} \widehat{t_{\mathrm{1j}}} & \widehat{t_{\mathrm{2j}}} & \cdots & \widehat{t_{N_{\mathrm{u}}\mathrm{j}}} \end{pmatrix}^T.$$

Each delay profile was then standardized by subtracting the mean and divided by standard deviation. By combining the standardized delay profiles for all time segments, we constructed a delay matrix $D$:

$$D = \begin{pmatrix} d_1 & d_2 & \cdots & d_{N_{segs}} \end{pmatrix},$$

where $N_{segs}$ is the number of time segments.

We then applied singular value decomposition (SVD) to the delay matrix $D$:

$$\mathrm{D} = \mathrm{U}\Sigma\mathrm{V}^{\mathrm{T}},$$

where $U, V$ corresponds to the left- and right-singular vectors, and $\Sigma$ contains singular values. The principal delay profile $u^*$ was defined as the first column of $U$ by which the maximum variance was explained. Therefore, the principal delay profile analysis identified a direction representing the temporal order of sequential activations across all the recorded neurons. The principal delay profile is unitless. We thus rescaled it to the unit of seconds based on the averaged cascade pattern (4).

#### Replays

The hippocampal replay was generally observed at a timescale of hundreds of milliseconds. To detect such fine-scale event, a replay detection algorithm was developed based on neural spiking activity binned by the 50ms time bins. Specifically, we applied the delay-profile decomposition method with the global mean signal low-pass filtered at 5Hz. The resting stationary periods were divided into fine-scale time segments $C_{\text{fine}}$ based on troughs of the filtered global signal. To detect the replays of the CA1 time-selective neurons, we computed the Spearman's rank correlation between the time fields of the CA1 time-selective neurons $\{TF_i^*\}_{i\in U_{CA1-T}}$ and the delay profile of these neurons at each fine-scale time segment. A replay event was identified if the absolute value of this correlation was larger than a pre-defined threshold. Therefore, the set of replays can be mathematically described as

$$\mathrm{R}_{\mathrm{CA1-T}} = \left\{ j \in C_{\text{fine}} \mid \left|\mathrm{corr}\left(\{TF_i^*\}_{i\in U_{CA1-T}}, \{d_{ij}\}_{i\in U_{CA1-T}}\right)\right| > r_{thr} \right\},$$

where $r_{thr}$ is the threshold that corresponds to the 99% significance level (Fisher Z-transformation P<0.01). The number of replays then was $|R_{CA1-T}|$.

We also considered three other cell ensembles as controls for comparison, the VIS-time-selective neurons $U_{\text{VIS-T}}$, the CA1-non-time-selective neurons $U_{\text{CA1-nT}}$ and the CA1-time-selective neurons for shuffled movies $U_{\text{CA1-T}^s}$, and their set of replays can be defined similarly:

$$R_{VIS-T} = \left\{ j \in C_{\text{fine}} \mid \left|\mathrm{corr}\left(\{TF_i^*\}_{i\in U_{VIS-T}}, \{d_{ij}\}_{i\in U_{VIS-T}}\right)\right| > r_{thr} \right\},$$

$$R_{CA1-\mathrm{n}T} = \left\{ j \in C_{\text{fine}} \mid \left|\mathrm{corr}\left(\{TF_i^*\}_{i\in U_{CA1-nT}}, \{d_{ij}\}_{i\in U_{CA1-nT}}\right)\right| > r_{thr} \right\},$$

$$R_{CA1-T^s} = \left\{ j \in C_{\text{fine}} \mid \left|\mathrm{corr}\left(\{TF_i^*\}_{i\in U_{CA1-T^s}}, \{d_{ij}\}_{i\in U_{CA1-T^s}}\right)\right| > r_{thr} \right\}.$$

For each of these neuron groups, we randomly shuffled the time fields within the group and then identified and counted the number of replays that occurred in the resting period. We repeated the procedure 500 times and constructed a null distribution for the number of replays occurred by chance. If the number of real replays is significantly larger than what would be expected from the null distribution, it suggests that such sequential replays occurred not by chance during the resting stationary periods. We approximated the null distribution with Gaussian and derived z-score to quantify how significantly the number of real replays differ from the null distribution.

The replay often comes in two directions, forward and reverse (6). We also classified the replays into forward and reverse based on the sign of the correlation between the time field and its resting-state delay profiles. Take CA1-time-selective neurons as an example, its forward replay is defined as

$$R_{CA1-T}^f = \left\{ j \in C_{\text{fine}} \mid \mathrm{corr}\left(\{TF_i^*\}_{i\in U_{CA1-T}}, \{d_{ij}\}_{i\in U_{CA1-T}}\right) > r_{thr} \right\},$$

whereas its reverse replay is defined as

$$R_{CA1-T}^r = \left\{ j \in C_{\text{fine}} \mid \mathrm{corr}\left(\{TF_i^*\}_{i\in U_{CA1-T}}, \{d_{ij}\}_{i\in U_{CA1-T}}\right) < -r_{thr} \right\}.$$

### Ripple detection

Hippocampal sharp waves ripples (SWRs) are short-lived fast oscillations (110-200Hz) observed in the LFP of the hippocampal recording sites. We adopted an offline ripple detection method (4, 7) to identify ripple events using the LFP signal recorded from the hippocampal CA1 region. The LFP was originally recorded at 2.5k Hz. In the Visual Coding dataset, it was later down sampled to 1.25K Hz. We performed ripple detection individually on the LFP from each CA1 recording site (channel) and the detected ripples are robust and largely overlapped across the channels. To integrate the detection results from different channels, we considered a detected ripple event valid only when it was detected in more than 40% CA1 channels. For verification, we computed the spectrogram with a sliding window of 20ms for each ripple event including a surrounding time window of 960ms. The spectrogram was normalized by subtracting the baseline power level in the first 100ms and then averaged across ripples to get the mean spectrogram. We similarly obtained the mean LFP pattern of ripple events by averaging the LFP signal over the surrounding 960ms time window of all ripple events (4).

### Slow coarse-scale cascades

Recently, it was found that the neural activation organized as spiking cascades of 5-15 seconds during immobile rest (4). In this paper, we termed such cascade as slow cascade so to differentiate it from the finer-scale micro-cascade. We followed the definition of the slow cascade in its original paper (4). Specifically, to detect the seconds-scale brain events, we applied the delay-profile decomposition method to the neural spiking data binned with 200ms time bins, with the global mean spiking rate being low-pass filtered at 0.5Hz. We thus obtained the principal delay profiles $u^*_{\text{coarse}}$ and a delay profile matrix $D$ where the columns are the delay profile of the coarse-scale time segments $C_{\text{coarse}}$. We further introduced a sequential score and assigned it to each time segment for identifying the slow cascades. The sequential score for the time segment $T_j$ is defined as

$$Seq(T_j) = \frac{\text{corr}(u^*_{\text{coarse}}, d_j)}{s_{thr}},$$

where $d_j$ is the delay profile of $T_j$, and $s_{thr}$ is the correlation value corresponding to the significance level of $p = 0.001$ (Fisher's Z-transform). Thus, the $|Seq(T_j)| > 1$ would suggest the significant sequential activations along or opposite to the principal delay profile direction. The slow cascades, denoted as $\Omega$, can be identified from the set of time segments $C_{\text{coarse}}$ by comparing the sequential score with 1:

$$\Omega = \{j \in C_{\text{coarse}} \mid Seq(T_j) > 1\}.$$

We followed the same procedure used by the previous study (4) to define two groups of neurons at the two ends of the principal delay profile, namely positive-delay neurons and negative-delay neurons. The positive-delay neurons were defined as those neurons with significant positive principal delay values in the cascades $\Omega$ whereas the negative-delay neurons are those with significant negative principal delay values ($p < 0.001$, one-sample t-test).

The slow cascade features a sharp increase in the spiking activity of positive-delay neurons in the middle, we thus identified and defined these time points as the positive-delay neuron onset. We also investigated how neuron population activity of each brain region was modulated around these time points at the slow cascades. For each brain region containing more than 100 recorded neurons, we computed its mean neural spiking activity at the slow cascades by aligning to the positive-delay neuron onsets.

To understand how the replays were distributed in the slow cascade cycle, we constructed an 8-second time window centering the positive-delay neuron onset of each slow cascade in Ω. A time course of replay events was constructed with value 1 for the existence of the replay and 0 for the absence:

$$r(t) = \begin{cases} 1 & t \in R_{CA1-T} \\ 0 & o.w. \end{cases}$$

We then averaged the replay time course $r(t)$ across the time windows of slow cascades to obtain the replay probability across the slow cascade cycle. As a control, we randomly shuffled the replay events sequence and computed the probability across the cascade cycle in the same manner. We then normalized the real probability curve by the mean probability from the control condition to quantify the relative possibility of the replay events.

**Micro-cascades**

Close inspection of spontaneous spiking data at the higher temporal resolution (time bin = 50 ms) suggested the existence of micro-cascade of hundreds of milliseconds. They showed similar sequential activations as the slow cascades of seconds but are much shorter and with very brief activation of the positive-delay neurons. Since they share a similar timescale as the replay events, we detected them from the fine-scale time segments $C_{\text{fine}}$ using the delay profile decomposition method described above. Similar to the slow cascade identification, we computed a fine-scale delay profile for each segment in $C_{\text{fine}}$ but then correlated it with the coarse-scale principal delay profiles. The resulting sequential scores quantify the existence of sequential activations of all the recorded neurons along this principal delay profile direction. The micro-cascades were bi-directional, either in a direction from the negative-delay neurons to the positive-delay neurons (N-P direction), or reversely, from the positive-delay neurons to the negative-delay neurons (P-N direction). Therefore, we defined the N-P cascade as

$$\Theta_{\text{N-P}} = \{\text{j} \in C_{\text{fine}} \mid Seq(T_j) > 1\}$$

whereas P-N cascade as

$$\Theta_{\text{P-N}} = \{\text{j} \in C_{\text{fine}} \mid Seq(T_j) < -1\}.$$

We obtained mean micro-cascade patterns by averaging the neural spiking activities over time windows aligned to the global spiking peaks of identified micro-cascades. We observed that the ripple events were aligned to the positive-delay neuron onset within the micro-cascades. Therefore, to investigate how ripple and replay events are modulated by the micro-cascade, we constructed a time window of 4 seconds centered at the positive-delay neuron onset of each micro-cascade and computed their normalized probability of occurrence over this time window. We also summarized the distribution of sequential scores for both forward and reverse replay separately. Finally, we performed statistical analyses on the relationship between (forward / reverse) replays and (P-N / N-P) micro-cascade. For each replay, we categorized it based on its sequential score into 3 classes, P-N micro-cascade, N-P micro-cascade, and non-micro-cascade. Then, the proportion of the (forward / reverse) replays categorized into different micro-cascade classes can be computed. To measure the significance, for each case, e.g., proportion of forward replay cooccurring with P-N micro-cascade, we constructed null distributions by computing the proportion of randomly shuffled replay sequence with 500 times repeat. We approximated the null distribution with Gaussian and derived the z-score to quantify the significance. It suggested that there is significant modulation between the micro-cascade and replay events (**Fig. S6C and S6D**).

The principal delay profile of fine-scale time segments $C_{\text{fine}}$, $u^{*}_{fine}$ was also derived by SVD. We rescaled it to the unit of seconds based on the averaged spiking pattern of N-P micro-cascades. We then correlated the fine-scale principal delay profile with the coarse-scale principal delay profile.

The fast-transitioning point of the slow cascades (i.e., around the positive-delay neuron onset) was often identified as a N-P micro-cascade. To verify the effect of other off-center micro-cascades, we excluded the micro-cascade occurring the positive-delay neurons onset of the slow cascades. Then the same analysis was performed on these off-center micro-cascades to investigate their relationship with ripples and replays (**Fig. S7**).
